# EPIGENE: genome wide transcription unit annotation using a multivariate probabilistic model of histone modifications

**DOI:** 10.1101/2019.12.17.878454

**Authors:** Anshupa Sahu, Na Li, Ilona Dunkel, Ho-Ryun Chung

## Abstract

**Background:** Understanding transcriptome is critical for explaining functional as well as regulatory roles of genomic regions. Current methods for the identification of transcription unit (TU) uses RNA-seq which, however, requires large quantities of mRNA limiting the identification of inherently unstable TUs e.g. for miRNA precursors. This problem can be resolved by chromatin based approaches due to a correlation between histone modifications and transcription.

**Results:** Here we introduce EPIGENE, a novel chromatin segmentation method for the identification of active TUs using transcription associated histone modifications. Unlike existing chromatin segmentation approaches, EPIGENE uses a constrained, semi-supervised multivariate hidden markov model (HMM) that models the observed combination of histone modifications using a product of independent Bernoulli random variables, to identify active TUs. Our results show that EPIGENE can identify genome-wide TUs unbiasedly. EPIGENE predicted TUs showed an enrichment of RNA Polymerase II in transcription start site and gene body indicating that they have been transcribed. Comprehensive validation with existing annotations revealed that 93% of EPIGENE TUs can be explained by existing gene annotations and 5% of EPIGENE TUs in HepG2 can be explained by microRNA annotations. EPIGENE outperforms existing RNA-Seq based approaches in TU prediction precision across human cell lines. Finally, we identify 381 novel TUs in K562 and 43 novel cell-specific TUs all of which are supported by RNA Polymerase II data.

**Conclusions:** We demonstrate the applicability of HMM to identify genome-wide active TUs and provides valuable information about unannotated TUs. EPIGENE is an open-source method and is freely available at: https://github.com/imbeLab/EPIGENE.

## 1. Background

Transcription unit (TU) represents the transcribed regions of genome which generates protein-coding genes as well as regulatory non-coding RNAs like microRNA. Accurate identification of TUs is important to better understand the transcriptomic landscape of the genome. With the rapid development of low‐cost high‐throughput sequencing technologies, RNA sequencing (RNA-seq) has become the major tool for genome‐wide TU identification. As a result, popular TU prediction tools such as AUGUSTUS [1], Cufflinks [2], StringTie [3], Oases [4] use RNA-seq data. Though RNA-seq based TU prediction can be considered the state-of-the-art method to annotate the genome, its main drawback lies in the dependence on relatively high quantities of target RNAs. This is problematic for accurate identification of inherently unstable TUs like primary miRNA. This shortcoming of RNA-Seq can be partly alleviated by chromatin-based approaches [5,6], due to the association between histone modifications and transcription.

Eukaryotic DNA is tightly packaged into macromolecular complex called chromatin, which consists of repeating units of 147 DNA base pairs (bp) wrapped around an octamer of four histones H2A, H2B, H3, and H4 called the nucleosome. Post-translational modifications (PTM) to histones in the form of acetylation, methylation, phosphorylation and ubiquitination, play an important role in the transcriptional process. These PTMs are added, read and removed by so called writers, readers and erasers. In this way nucleosomes serve as signalling platforms [7] that enable the localized activity of chromatin signalling networks partaking in transcription and other chromatin-related processes [8]. Indeed, it has been shown that histone modifications are correlated to the transcriptional status of chromatin [9,10]. For example, H3K4me3 and H3K36me3 are positively correlated with transcription initiation [11,12] and elongation [13] and are considered as transcription activation marks, whereas H3K9me3 and H3K27me3 [11,14], are considered as repressive marks as they are commonly found in repressed regions. Therefore, it is reasonable to assume that histone modifications profiles can be used to identify cell-type-specific TUs. Given a deluge of cell-type-specific epigenome data available through many consortia, such as ENCODE [15], NIH Roadmap Epigenomics [16], DEEP [17], Blueprint [18], CEEHRC [19] and IHEC [20], a highly robust TU annotation pipeline based on epigenome markers becomes feasible.

Currently many computational approaches such as ChromHMM [21], EpicSeg [22], chroModule [23], GenoSTAN [24] etc., have been developed that use histone modifications as an input to provide a genome annotation. These chromatin segmentation approaches use a variety of mathematical models with most prominent one being hidden markov models (HMM). HMMs are a powerful tool for chromatin state identification based on histone modifications, due to their assumption that a combination of histone modifications is generated by an underlying hidden chromatin state emitting a combination of histone modifications according to a particular probability distribution.

Based on the training, these HMMs can be classified as: (1) unsupervised (methods like ChromHMM, EpicSeg and GenoSTAN), that do not include prior biological information and require user to interpret and annotate the learned states based on existing knowledge about functional genomics. (2) supervised (methods like chroModule), that relies on a set of positive samples to train on and consequently yields predictions that reflect the properties of the training set. Although these approaches annotate genome modules such as promoter, enhancer, transcribed regions etc, they fail to identify active TUs as they do not constrain the chromatin state sequence to begin with a transcription start site (TSS) and end with a transcription termination site (TTS).

To address these shortcomings, we developed a semi-supervised HMM, EPIGENE (EPIgenomic GENE), which is trained on the combinatorial pattern of IHEC class I epigenomes (H3K27ac, H3K4me1, H3K4me3, H3K36me3, H3K27me3 and H3K9me3) that are indicative of active transcription to infer the hidden “transcription unit state”. The emission probabilities represent the probability of a histone mark occurring in a TU state and the transition probabilities capture the topology of TU states. The HMM comprises of TU states and background states. The transcription start site (TSS), exons (first, internal and last exon), introns (first, internal and last intron) and transcription termination site (TTS) are referred to as the TU states. As, every TU begins with a TSS state, proceeds through intragenic states like exon and intron and terminates with a TTS state, a background state can only be reached from a TTS state and a TSS state can only be reached from a background or TTS (in case of genes occurring in close proximity to each other) state.

In the forthcoming sections, we describe the method, validate the predicted EPIGENE transcription units with existing annotations, RNA-Seq and ChIP-seq evidence, compare the performance of EPIGENE to existing RNA-Seq based TU prediction methods within and across cell lines and show that EPIGENE outperforms state-of-art RNA-Seq based approaches in prediction resolution and precision. In summary, EPIGENE yields predictions with a high resolution and provides a pre-trained model that can robustly be applied across samples.

## 2. Results and discussion

### 2.1 Schematic overview of EPIGENE

EPIGENE uses a multivariate HMM (shown in Figure 1A (ii)), which allows the probabilistic modelling of the combinatorial presence and absence of multiple IHEC class I histone modifications. It receives a list of aligned ChIP and control reads for each histone modification, which are converted into presence or absence calls across the genome using normR (see Materials and Methods section 4.5). By default, TU states are analysed at 200 bp non-overlapping intervals called bins. The HMM comprises of 14 TU states and 3 background states where each transcription unit state captures individual elements of gene such as TSS, exons, introns and TTS. The transition probability of transcription unit states were trained in a supervised manner using GENCODE annotations [25] and their emission probabilities were trained on a highly confident set of GENCODE transcripts [25] which showed an enrichment for RNA Polymerase II in K562 cell line (see Materials and Methods section 4.7). The transition and emission probabilities of background states were trained in an unsupervised manner (see Materials and Methods section 4.7). The HMM outputs a vector where each bin is assigned to a TU or background state, which is further refined to obtain active TUs (see Figure 1B).

**Figure 1:**
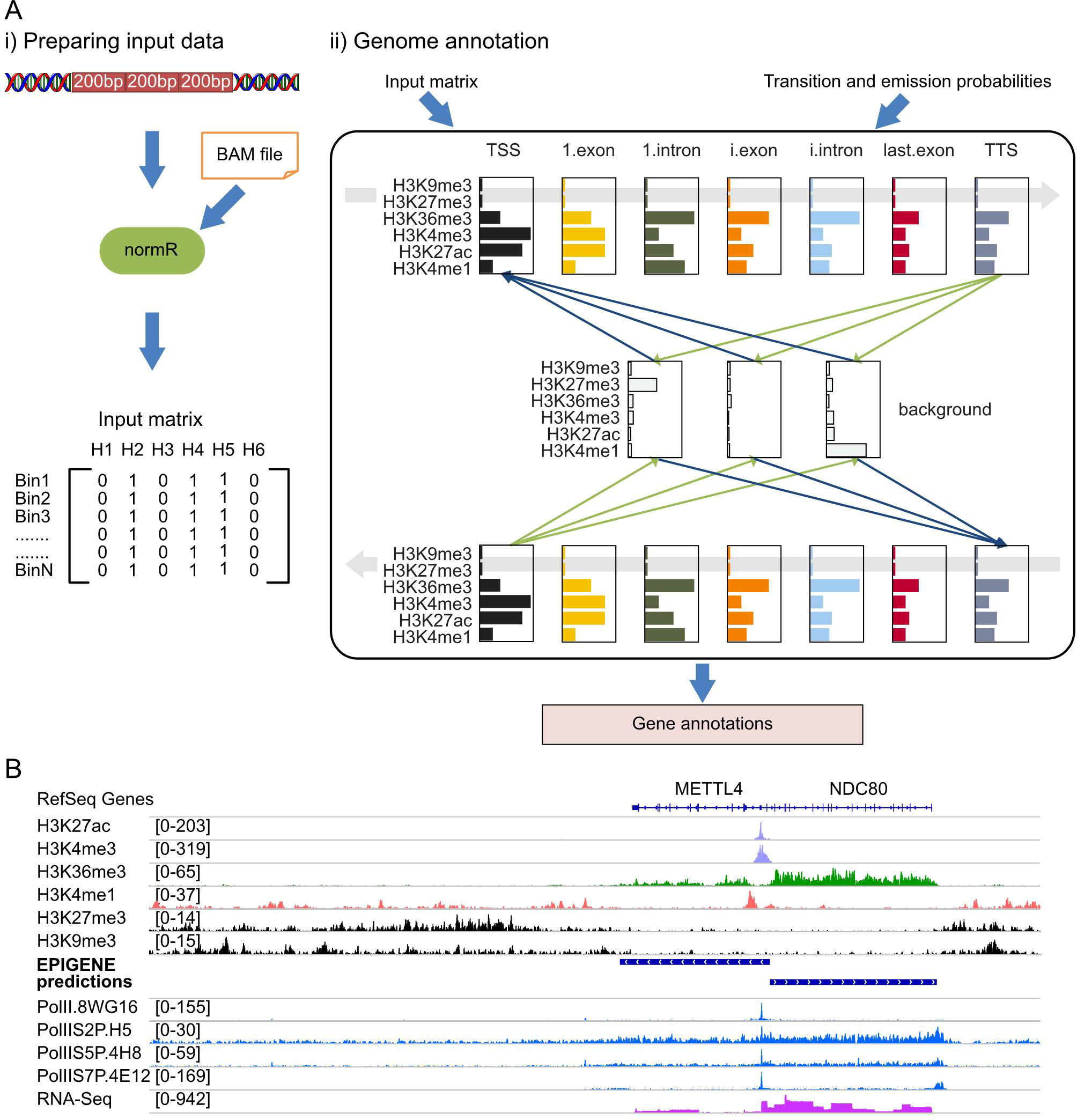
**A.** Schematic overview of EPIGENE framework. **B.** An example of EPIGENE prediction. EPIGENE predictions of METTL4 and NC80 gene, show an enrichment of H3K27ac and H3K4me3 at TSS (tracks shown in light violet), H3K36me3 in gene body (tracks shown in green), enhancer mark H3K4me1 few bps upstream or downstream of TSS (tracks shown in pink), RNA Polymerase II in TSS and gene body (tracks shown in blue). The predictions also show an absence of repression marks H3K27me3 and H3K9me3 (tracks shown in black). The corresponding RNA-Seq evidence in this genomic region can be seen in lower most track (track shown in dark pink)

### 2.2 Validation with existing gene annotations and RNA-Seq

We validate the predicted transcription units with existing gene annotations and RNA-Seq evidence, for this we combined the EPIGENE predictions (24,571 TUs) and RNA-Seq predictions that was obtained from Cufflinks (32,079 TUs) and StringTie (101,656 TUs; refer Table 2-4 in Supplementary file A1 for summary statistics) to generate a consensus TU set. This consensus TU set comprises of 24,874 TUs, which were then overlaid with GENCODE and CHESS gene annotation [25,26] (Figure 2). We find that 93% of EPIGENE TUs can be explained by existing gene annotations. We additionally identified 14,797 (11,584: annotated, 3213: unannotated) RNA-Seq-exclusive TUs and 1304 (718: annotated, 586: unannotated) EPIGENE-exclusive TUs, of which 65% of EPIGENE and 31% of RNA-Seq unannotated predictions show enrichment of RNA Polymerase II. Additional details about RNA Polymerase II enrichment in the consensus TU set can be seen in Supplementary table S1.

**Figure 2:**
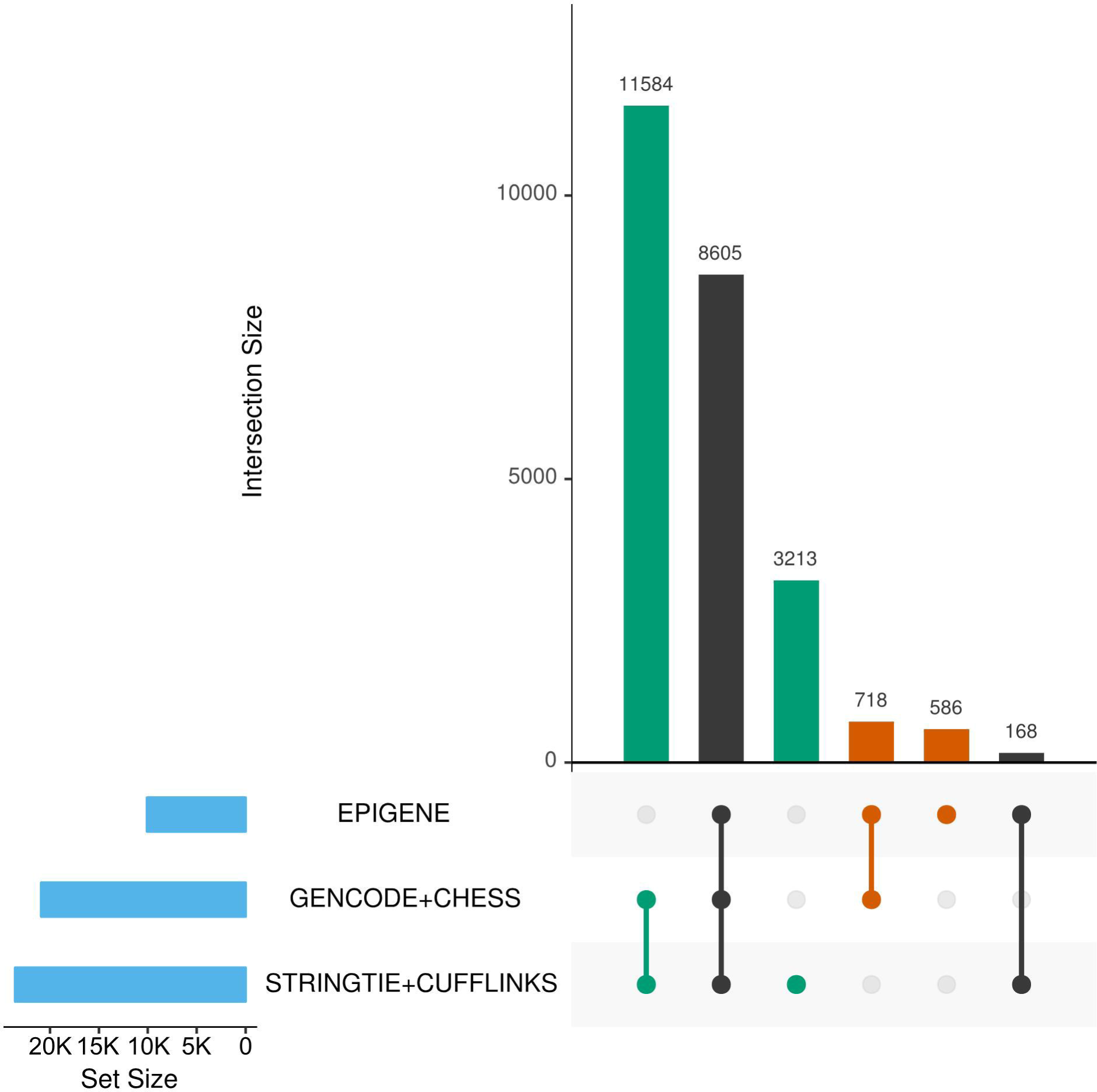
Overlap of EPIGENE predictions with existing gene annotations and RNA-Seq based predictions

### 2.3 Histone modifications and RNA Polymerase II occupancy

The correctness of predicted transcription units was estimated in K562, due to the availability of matched RNA Polymerase II and RNA-Seq profiles. We predicted 24,571 TUs in K562 cell line, majority of which showed typical gene characteristics, with high enrichment of H3K27ac, H3K4me3 and H3K36me3 in TSS and gene bodies (Figure 3A).

**Figure 3:**
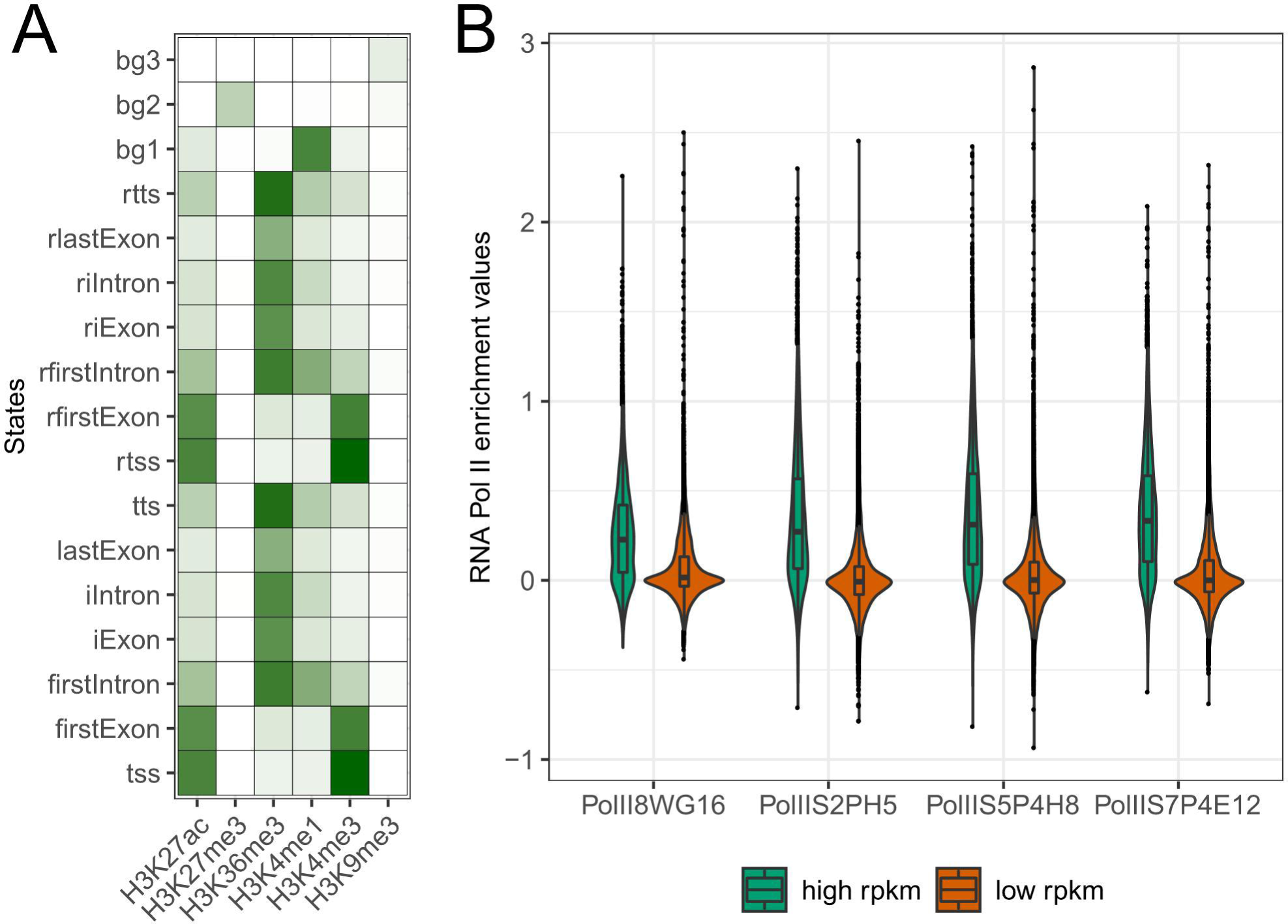
Correctness of EPIGENE predictions. **A.** EPIGENE estimated parameters for K562 using 17 chromatin states, ranging from 0 (white) to 1 (dark green). **B.** Distribution of RNA Polymerase enrichment score in EPIGENE predictions, predictions are divided as: high RPKM (RPKM >= upper quartile) and low RPKM (RPKM < upper quartile) based on RNA-Seq evidence in predicted transcripts

It is already known that eukaryotic transcription is regulated by phosphorylation of RNA Polymerase II carboxy-terminal domain in serine 2, 5 and 7. The signal for serine 5 and 7 is strong at promoter region where as signal for serine 2 and 5 phosphorylation is strong at actively transcribing regions [27]. Therefore, we incorporated RNA Polymerase II evidence in all the forthcoming analyses. Genome wide RNA Polymerase II profile for K562 cell line was obtained using four antibodies that capture RNA Polymerase II signal at transcription initiation and gene bodies. The enrichment of RNA Polymerase II in predicted TUs was computed using normR [28] (see Materials and Methods section 4.5). The predicted TUs were classified as: high and low RPKM based on mRNA levels (threshold = upper quartile). Figure 3B shows the distribution of RNA Polymerase II enrichment in both the classes of predicted TUs. We observe a significant proportion of predicted TUs (78%) show a positive enrichment score indicating the biological correctness of our predictions. We also come across 24 unannotated TUs that report an enrichment score above 0.5 but have a reduced or no RNA-Seq evidence.

### 2.4 Method comparison

Currently multiple approaches exist for predicting TU that rely on RNA-Seq evidence. We compare the performance of EPIGENE with two existing RNA-Seq based transcript prediction approaches, Cufflinks and StringTie, both of which are known to predict novel TUs in addition to annotated TUs. The method comparison was performed in two stages: within cell type and cross cell type comparison using RNA Polymerase II enrichment as performance indicator (see Materials and methods section 4.8). The confusion matrix defining the true positives (TP), true negatives (TN), false positives (FP) and false negatives (FN) can be seen in Figure 4A.

**Figure 4:**
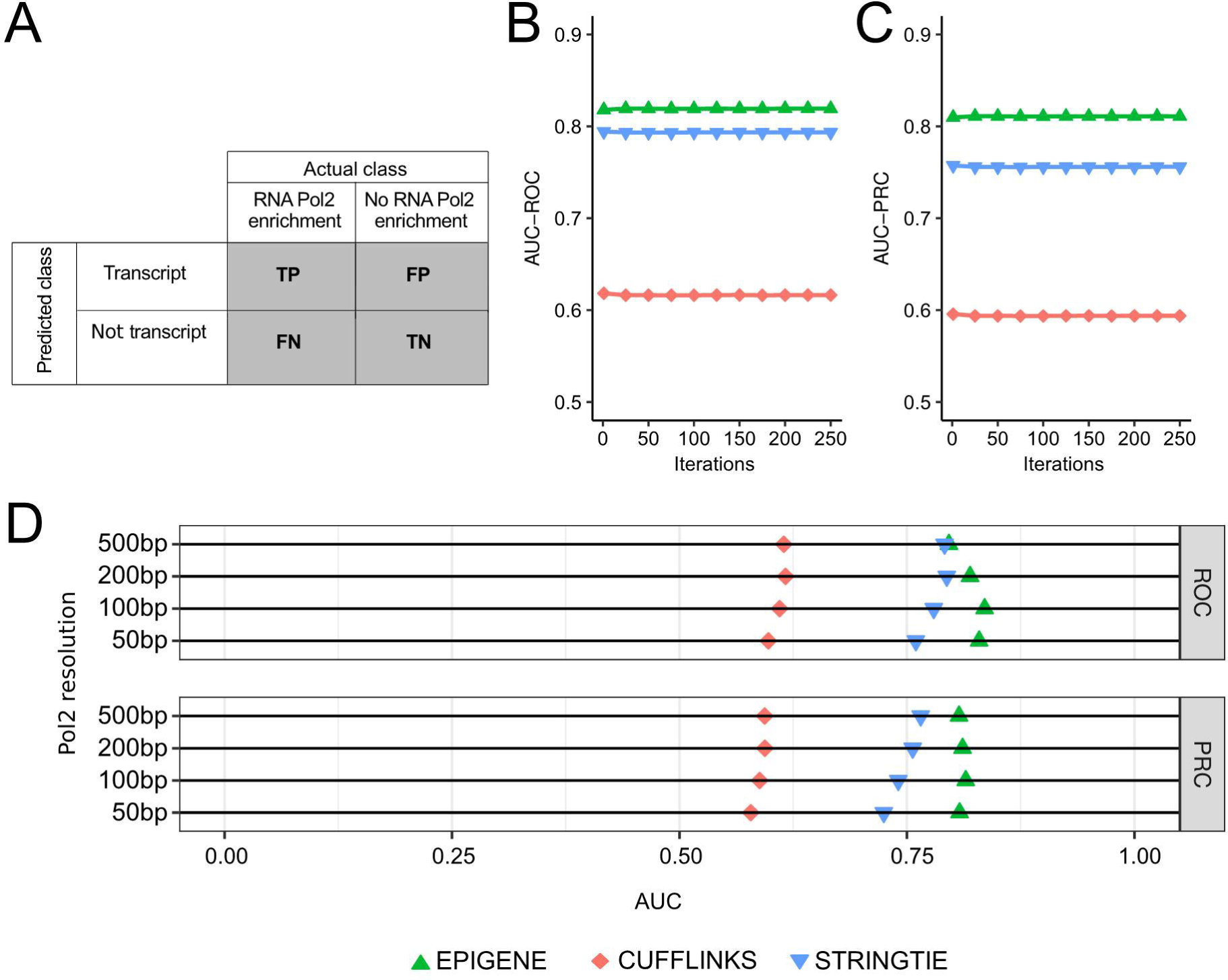
Performance of EPIGENE compared to existing RNA-Seq based transcription unit annotation methods: Cufflinks and StringTie. **A.** Contingency matrix used for method comparison. **B.** Receiver Operating Characteristic curve **C.** Precision-Recall curve. **D.** Area under ROC and PRC curve for varying RNA Polymerase II resolution for EPIGENE, Cufflinks and StringTie

#### 2.4.1 Within cell type comparison

For this comparison, we use the ChIP-seq profile of RNA Polymerase II in K562 cell line that was obtained using PolIIS5P4H8 antibody, due to its ability to identify RNA Polymerase II occupancy in TSS and actively transcribed regions.

As, evident from Figure 4B and 4C, EPIGENE outperforms both the RNA-Seq based approaches and reports a higher AUC (PRC: 0.81, ROC: 0.82) in both the curves compared to Cufflinks (PRC: 0.59, ROC: 0.64) and StringTie (PRC: 0.75, ROC: 0.79). The above analysis was repeated for varying resolutions (50,100 and 500 bp); the AUC reported for varying resolution can be seen in Figure 4D. As observed in the figure, Cufflinks achieve a lower AUC compared to StringTie and EPIGENE, which is likely due to the usage of the RABT assembler which results in large number of false positives [29].

EPIGENE reports a higher AUC than StringTie for varying RNA Polymerase II resolutions, this can be due to (1) the usage of RNA Polymerase II enrichment as a performance measure might lead to a ChIP-seq biasness towards EPIGENE, which is also a ChIP-seq based approach. This results in more true positives compared to RNA-Seq based approaches, or (2) RNA-mapping artefacts that results in more false positives than EPIGENE. Therefore, we examined the precision, sensitivity and specificity values for EPIGENE, Cufflinks and StringTie and found that the increased AUC for EPIGENE is due to spurious read mappings of RNA-Seq that results in higher false positives in StringTie and Cufflinks. Figure S2 (included in Supplementary file A1) shows an example of Cufflinks and StringTie TU that was identified due to spurious read mapping. This TU exactly overlaps with a repetitive sequence that occurs in four chromosomes (chromosome 1, 5, 6, X).

#### 2.4.2 Cross cell type comparison

In order to evaluate the performance of EPIGENE across cell types, we applied K562-trained models to samples from different cell types. We compared the approaches on three different datasets provided by the ENCODE [15] and DEEP [17,30] consortium:

1. IMR90: lung fibroblast cells with 6 histone modifications,1 RNA Polymerase II, two control experiments (one each for RNA Polymerase II and histone modifications) and one RNA-Seq obtained from ENCODE,
2. HepG2_1 and HepG2_2: hepatocellular carcinoma with 6 histone modifications, one control experiment and one RNA-Seq obtained from DEEP where two replicates per histone modification and RNA-Seq are available, RNA Polymerase II ChIP and control experiments obtained from ENCODE.

As shown in Figure 5 A, B and C, K562-trained EPIGENE models consistently achieve a higher prediction accuracy, outperforming Cufflinks and StringTie.

**Figure 5:**
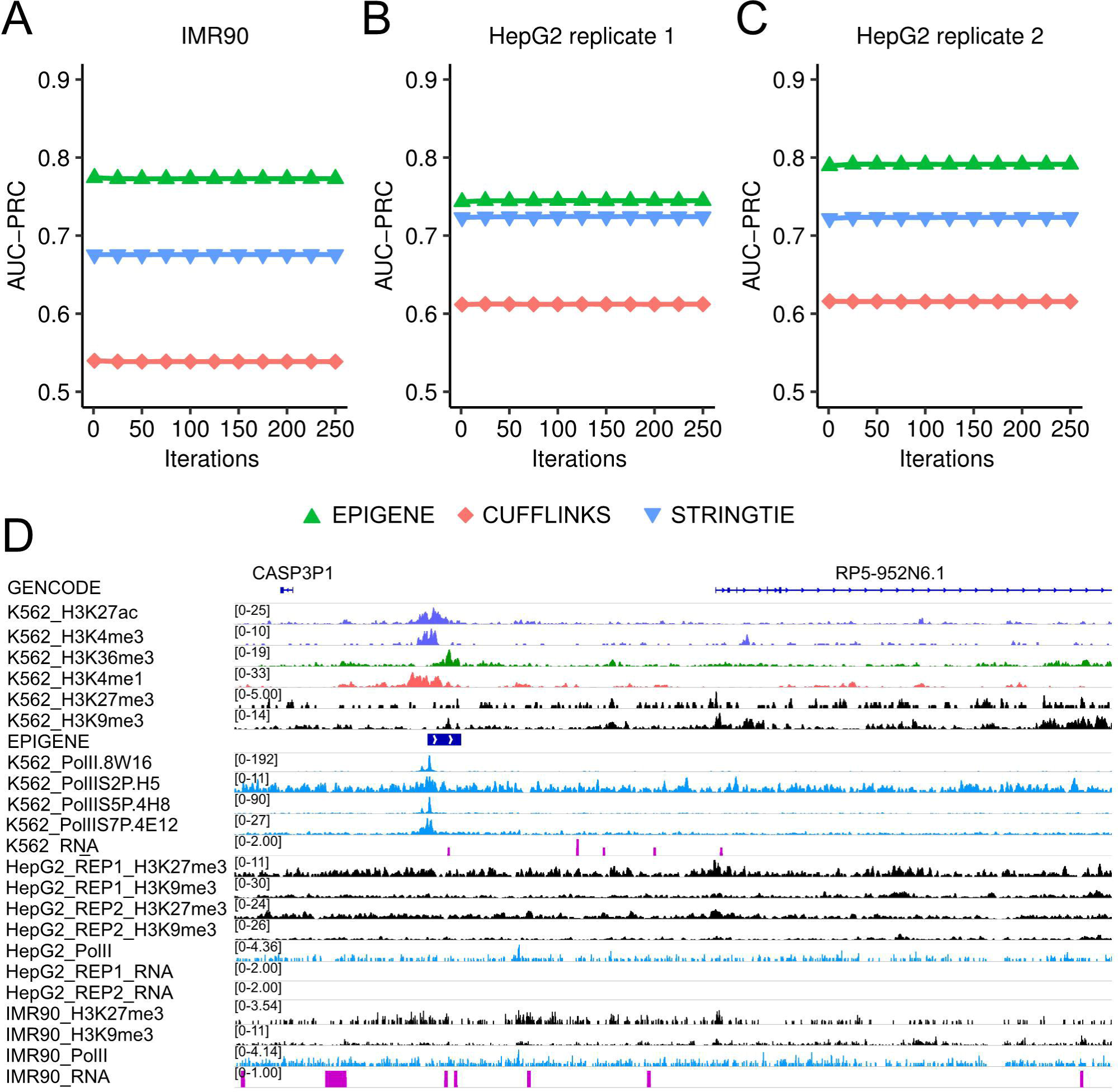
**A-C.** Performance of K562-trained EPIGENE models, Cufflinks and StringTie across cell lines. **D.** Example of EPIGENE predicted TU that lacks RNA-Seq evidence (tracks shown in dark pink). The TU was predicted to be active in K562 but not in HepG2 and IMR90, and is located between pseudogene CASP3P1 and lncRNA RP5-952N6.1. The TU shows an enrichment of H3K27ac and H3K4me3 at TSS (tracks shown in light violet), H3K36me3 in gene body (tracks shown in green), enhancer mark H3K4me1 few bps upstream of TSS (tracks shown in pink), K562 RNA Polymerase II in TSS and gene body (tracks shown in blue). The TU also show an absence of repression marks H3K27me3 and H3K9me3 in K562 (tracks shown in black). We additionally observe the enrichment of repression mark in H3K27me3 in HepG2 and IMR90 indicating that the region is repressed in both these cell lines

### 2.5 EPIGENE identifies transcription units with negligible RNA-Seq evidence

Previous analyses (see section 2.3 and 2.4) indicated the presence of transcription units with RNA Polymerase II evidence and reduced or no RNA-Seq evidence. Here we evaluate these transcription units within and across cell lines by: (1) identifying cell-type specific transcription units that show gene characteristics but lack RNA-Seq evidence, and (2) looking for the presence of microRNAs that were not identified by RNA-Seq.

#### 2.5.1 EPIGENE identifies cell-type specific transcription units

We create a consensus set of transcription units by overlaying the EPIGENE predictions from K562, HepG2 and IMR90. This consensus TU set comprised of 18,248 TUs, of which ~78% showed an enrichment for RNA-Polymerase II. We identified 10,233 differential TU, of which 8047 were exclusive to cell lines (K562: 4247, IMR90: 2545, HepG2: 1255; see Figure S3 in Supplementary file A1). We additionally identified 43 highly confident cell-specific TUs (K562: 24, IMR90: 17, HepG2: 2; additional details in Supplementary table S2) which lacked RNA-Seq evidence but showed typical characteristics of a TU, with RNA Polymerase II enrichment at TSS and transcribing regions, H3K4me3 and H3K27ac enrichment at the TSS and H3K36me3 enrichment in gene body. An example of one such TU can be seen in Figure 5D.

#### 2.5.2 Identifying microRNAs that lack RNA-Seq evidence

MicroRNAs are small (~22 bp), evolutionally conserved non-coding RNAs [31,32] derived from large primary microRNAs (pri-miRNA), that are processed to ~70 bp precursors (pre-miRNA) and consequently to their mature form by endonucleases [33,34]. They regulate various fundamental biological processes such as development, differentiation or apoptosis by means of post-transcriptional regulation of target genes via gene silencing [35,36] and are involved in human diseases [37]. Due to the unstable nature of primary microRNA, traditional identification approaches relying on RNA-Seq are challenging. Here, we investigate the presence of primary microRNA that lack RNA-Seq evidence across cell lines. We create a consensus TU set (used in section 2.2) for individual cell lines (K562, HepG2 and IMR90) and overlaid them with miRbase annotations [38] to obtain potential primary microRNA TUs. We identified 655 EPIGENE TUs (5% of total EPIGENE TUs common in both replicates) that can be explained by miRbase annotations. We observe that majority of these are supported by RNA-Seq and Polymerase II evidence (Figure 6A and Figure S4 Supplementary file A1). We additionally identify 2 primary microRNA TUs in HepG2 cell line, which showed an enrichment for H3K4me3 in promoters, H3K36me3 in gene body and RNA Polymerase II in TSS and transcribing regions; and lacked RNA-Seq evidence. One of these transcription units overlaps with a microRNA cluster located between RP-11738B7.1 (lincRNA) and NRF1 gene (see Figure 6B).

**Figure 6:**
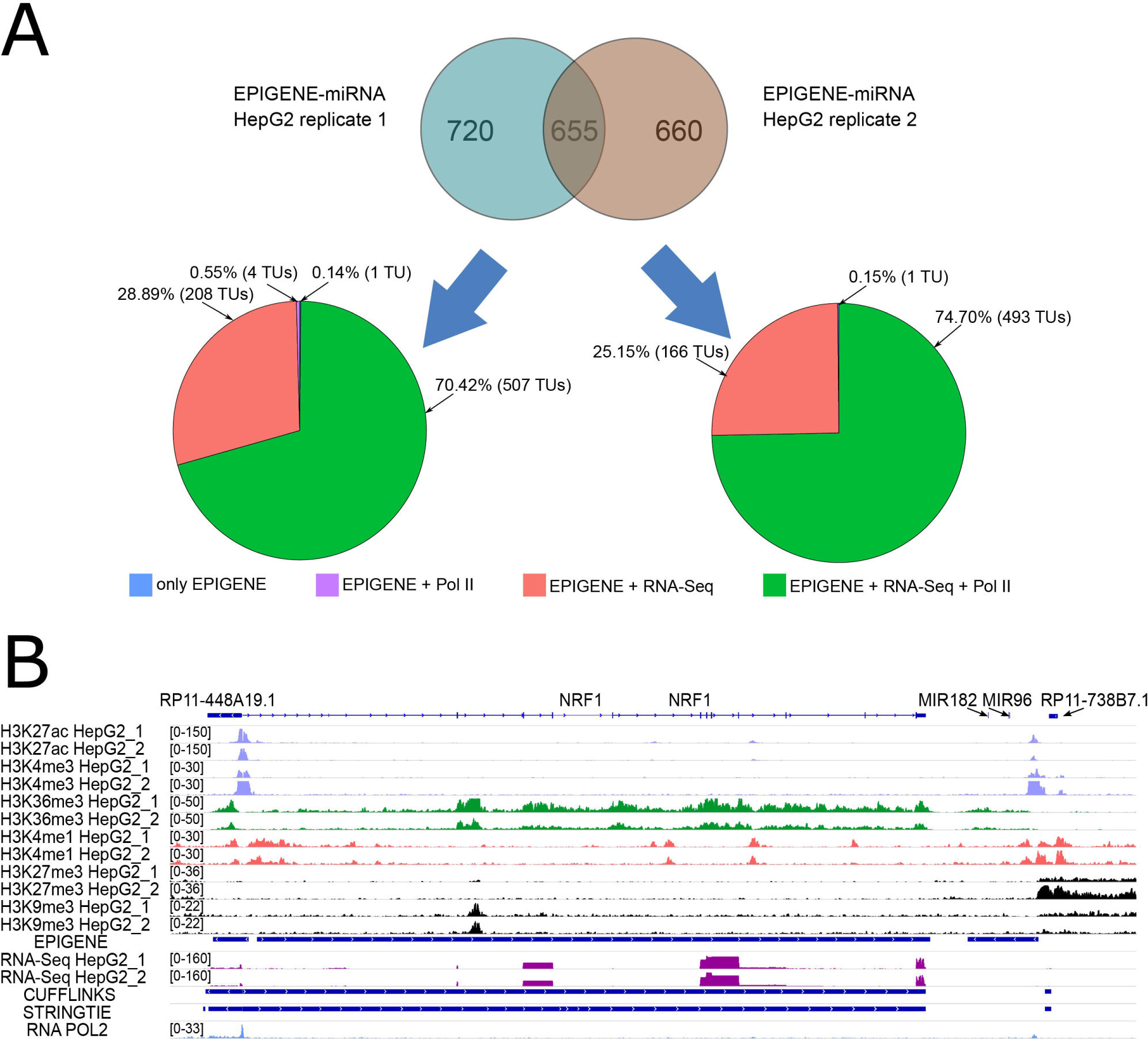
**A.** Overview of potential primary miRNAs predicted by EPIGENE in HepG2. **B.** Example of a TU overlapping a microRNA cluster was predicted by EPIGENE in HepG2 cell line. This region is located between lincRNA RP11-738B7.1 and gene NRF1 which was identified as a key player in maintaining cellular homeostasis and organ integrity [46]. The TU shows an enrichment of H3K27ac and H3K4me3 at TSS (tracks shown in light violet), H3K36me3 in gene body (tracks shown in green), enhancer mark H3K4me1 few bps upstream and downstream of TSS (tracks shown in pink), RNA Polymerase II in TSS (tracks shown in blue). The predictions also show an absence of repression marks H3K27me3 and H3K9me3 (tracks shown in black) and RNA-Seq evidence (tracks shown in dark pink).

### 2.6 Discussion

In this work, we introduced EPIGENE, a semi-supervised HMM that identifies active TUs using histone modifications. EPIGENE comprises of TU and background sub-models. The TU sub-model was trained in a supervised manner on predefined training sets, while the background was trained in an unsupervised manner. This semi-supervised approach captures (1) the biological topology of active TUs, and (2) probability of occurrence of histone modifications in different parts of a TU.

We first showed that majority of the predicted TUs can be explained by existing gene annotations, histone modifications and RNA Polymerase II. A quantitative comparison with RNA-Seq reveals the presence of TUs with RNA-Polymerase II enrichment but negligible RNA-Seq evidence. Considering RNA-Polymerase II as true transcription indicator, we compared the performance of EPIGENE with two RNA-Seq based approaches Cufflinks and StringTie. Based solely on the AUC of PRC and ROC curve as performance measure, EPIGENE achieves a superior performance than RNA-Seq based approaches. We further showed that EPIGENE can be reliably applied across different cell lines without the need for re-training and accomplishes a superior performance than RNA-seq based approaches.

We examine other performance scores like precision, sensitivity and specificity values, and observe that the high AUC of EPIGENE is due to RNA Seq mapping artefacts that result in high number of false positive in Cufflinks and StringTie. We further evaluate the presence of differentially identified TUs in K562, HepG2 and IMR90 cell line that lack RNA-Seq evidence. The results suggest the presence of cell line exclusive transcripts that lack RNA-Seq evidence. We additionally identify microRNAs that lack RNA-Seq evidence due to their labile nature. All of the aforementioned TUs show an enrichment of RNA Polymerase II in TSS and gene body indicating that they have been transcribed.

It is important to note that EPIGENE does not differentiate between functional and non-functional units of a TU (exons and introns) as the association between histone modifications and alternative splicing is yet to be elucidated [39]. However, EPIGENE identifies active TUs with greater precision as shown in section 2.4 and in the example regions presented in this work. The accuracy of EPIGENE predictions depends on the sequencing depth of the input histone modifications, therefore, high quality ChIP-seq profiles of histone modifications should be used to obtain confident transcription unit annotation.

Altogether, the superior performance within and across cell lines, identification of TUs especially primary microRNAs lacking RNA-Seq evidence as well as interpretability makes EPIGENE a powerful tool for epigenome based gene annotation.

## 3. Conclusion

With increasing efforts in the direction of epigenetics, many consortia continue to provide high quality genome-wide maps of histone modifications but determining the genome-wide transcriptomic landscape using this data has remained unexplored so far. Extensive evaluations in this work demonstrated the superior accuracy of EPIGENE over existing transcript annotation methods based on true transcription indicators. EPIGENE framework is user-friendly and can be executed by solely providing binarized enrichments for ChIP-seq experiments, without the need to re-train the model parameters. The resulting transcript annotations are in good agreement with RNA-Polymerase II evidence and can be used to provide a cell specific, epigenome-based gene annotation.

## 4. Materials and methods

### 4.1 Library preparation of histone modifications ChIP-seq

For K562 cell line presented in this study, ChIP against six core histone modifications, H3K27ac, H3K27me3, H3K4me1, H3K4me3, H3K36me3 and H3K9me3, was performed. The sheared chromatin without antibody (input) served as control. 10 × 10^6^ K562 cells were cultured as recommended by ATCC. Chromatin immunoprecipitations were preformed using the Diagenode auto histone ChIP seq kit and libraries were made using microplex kits according to manufacturer’s instructions and 10 PCR cycles.

### 4.2 Library preparation of RNA Polymerase II ChIP-seq

K562 cells were cultured in IMDM (#21980Gibco) with 10% FBS and P/S. Cells at a concentration of 1.2mio/ml were fixed with 1% Formalin at 37°C for 8min. Nuclei were isolated with a douncer, chromatin concentration was measured and 750µg chromatin per CHIP was used. Samples were sonicated with Biorupter for 33 cycles (3x 11 cycles). Chromatin, antibodies (RNA Pol II Ser2P (H5), RNA Pol II Ser5P (4H8), RNA Pol II Ser7P (4E12) and PolII (8WG16)) and protein G beads were combined and rotated at 4°C. For elution 250µl elution buffer (1% SDS) was used and after reverse crosslinking DNA was isolated by Phenol Chloroform extraction and elute in 1xTE. Final concentration was measured by Qubit. Bioanalyzer was done to check fragment sizes.

### 4.3 Sequencing and processing of ChIP-seq data

Sequencing for RNA-Polymerase II and histone modifcations was performed on an Illumina Highseq 2500 using a paired end 50-flow cell and version 3 chemistry. The resulting raw sequencing reads were aligned to the genome assembly “hs37d5” with STAR [40] and duplicates were marked using Picard tools [41]. We used *plotFingerprint* which is a part of deepTools [42] to access the quality metrics of for all ChIP-seq experiments.

### 4.4 Processing of RNA-Seq data

The raw reads from RNA-Seq experiments were downloaded from European Nucleotide Archive (SRR315336, SRR315337 for K562), European Genome Archive (EGAD00001002527 for HepG2) and ENCODE (ENCSR00CTQ for IMR90) and were aligned to the genome assembly “hs37d5” with STAR [40].

### 4.5 Binarization of enrichment levels

EPIGENE requires the enrichment values of IHEC class I histone modifications in a binarized data form or a "class matrix" to learn a transcription state model. This was done by partitioning the mappable regions of the genome of interest into non-overlapping sub-regions of the same size called bins. In the current setup, the transcription states are analysed at 200bp resolution, as it roughly corresponds to the size of a nucleosome and spacer region. Given the ChIP and input alignment files for each of the histone modifications, the class matrix for multivariate HMM is generated using the following approach:

1. *Obtaining read counts*: Read counts for all the bins is performed using *bamCount* method from R package *bamsignals* [43], with the following parameter settings: mapqual = 255, filteredFlag = 1024, paired.end = midpoint.
2. *Enrichment calling and binarization*: After having obtained the read counts, enrichment and binarization for each of the histone modification across all bins is computed using *enrichR* (binFilter = zero) and *getClasses* (fdr = 0.2) method from *normR* [28], which uses a negative binomial distribution to perform enrichment and binarization. This step yields the class matrix that serves as an input for the multivariate HMM.

### 4.6 The EPIGENE model

EPIGENE uses a multivariate HMM (shown in Figure 1A (ii)) to model the class matrix and identify active transcription units. Class matrix *C* is a m × n matrix, where, ***m*** = total number of 200 bp bins, and, ***n*** = number of histone modifications. Each entry *Cij* in the class matrix *C* corresponds to the binarized enrichment in i-th bin for the j-th histone modification. The model constitutes *k* number of hidden states (which is an input parameter of the algorithm), and each row of the class matrix corresponds to a hidden state. The emission probability vector for each hidden state corresponds to the probability with which each histone mark is found for that hidden state. The transition probabilities between the states enables the model to capture the position biases of gene states relative to each other. The emission probabilities of each state represents the probability with which each histone mark occurs in a state. Given this model, the algorithm does the following:

1. Initializes the emission, transition, and initial probabilities.
2. Fits the emission, transition, and initial probabilities using the Baum-Welch algorithm [44].
3. As we are concerned about the most probable sequence of active transcription unit, therefore, the sequence of hidden states is inferred using the Viterbi algorithm [45].

### 4.7 Training the model parameters

The transition and emission probabilities of the multivariate HMM are trained using GENCODE annotations with the following approach.

1. Bins overlapping gencode transcripts are identified and termed as gencode bins.
2. The gencode bins were categorized as TSS, TTS, 1st, internal and last exon and intron bins, and were subsequently grouped based on transcript IDs.
3. The coverage (in bp) of individual transcription unit component (i.e TSS, 1st exon, 1st intron etc) for each transcript is computed to generate the coverage list, where each entry of the coverage list contains the coverage information (in bp) for individual transcripts.
4. The transition probability of each “transcription unit state” was computed from the coverage list, and the missing probabilities from and to the “background state” are generated in an unsupervised manner.
5. We filtered the gencode transcripts to obtain transcripts that report an enrichment for RNA Polymerase II. This was done by clustering the binarized enrichment values of RNA Polymerase II in TSS and TTS bins of the transcripts and obtaining TSS and TTS bins that reports a high cluster mean for RNA Polymerase II. The emission probability of each “transcription unit state” was computed from class matrix and coverage of these transcripts (coverage computed from Step 2). The missing emission probabilities for the background states are trained in an unsupervised manner.

### 4.8 Performance evaluation

The performance of EPIGENE and RNA-Seq based transcript prediction approaches is evaluated using RNA Polymerase as performance indicator. This is done by removing assembly gaps in the genomic regions of interest and partitioning the remaining contigs into non-overlapping bins of 200 bps. The actual transcription status of each 200 bp bin was given by the observed binarized RNA Polymerase II enrichment in the bin and the predicted transcription status of the bin for method m, ***PT***_***m***_(***bin***) is given by:

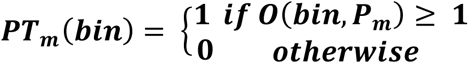

where, ***O***(***bin***, ***P***_***m***_) is the overlap between the bin and method m predictions ***P***_***m***_.

The predictions of EPIGENE and other RNA-Seq based approaches is evaluated by computing the area under curve for Precision-Recall (AUC-PRC) and Receiver Operating Characteristic curve (AUC-ROC) with primary focus on AUC-PRC. Considering a very high class imbalance i.e. 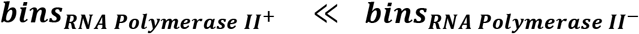, the AUC-PRC and AUC-ROC is computed using random sampling as:

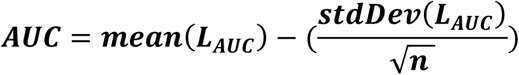

where, ***n*** is the sampling size or number of iterations and ***L***_***AUC***_ is the list of AUCs obtained for sampling size ***n***.

## Declarations

### Ethics approval and consent to participate

Not applicable

### Consent for publication

Not applicable

### Availability of data and material

Data for ChIP-seq experiments for K562 cell line are available via European Nucleotide Archive (PRJEB34999). Additional details about other ChIP-seq and RNA-Seq data used in this work can be found in the Supplementary file A1, Table 1. EPIGENE code is available at: https://github.com/imbeLab/EPIGENE.

### Authors’ contributions

The project was conceived by HC. AS performed all the analyses and wrote the manuscript with inputs from HC. NL performed the ChIP-seq for histone modifcations in K562. ID performed the ChIP-seq for RNA Polymerase II in K562.

## Acknowledgements

The authors would like to thank Clemens Thoelken for helpful comments on the manuscript. Many thanks to Sarah Kinkley, Anna Ramisch, Tobias Zehnder and Giuseppe Gallone from MPIMG for their valuable comments and inspiring discussions.

## List of supplementary files

| File name | File format | Title | Description of data |
| --- | --- | --- | --- |
| **Supplementary_table-S1** | .csv | RNA Polymerase II enrichment | RNA Polymerase II enrichment in consensus TU set |
| **Supplementary_table-S2** | .csv | Cell specific TUs | Additional details about cell specific TUs that lack RNA-Seq evidence |
| **Supplementary_file_A1** | .pdf | Data details and additional results | Details of datasets used and additional results |

