## Supplementary file A1 for "EPIGENE: genome wide transcription unit annotation using a multivariate probabilistic model of histone modifications"

### 1. Data

Table 1 lists the datasets used in this study. The RNA-Seq and ChIP-seq data were processed as mentioned in Section 4.3. The sequencing reads were binned in 200 bp bins and the raw ChIP read counts were normalized and binarized using normR. In this study we consider only uniquely mapped reads.

| Cell line | Residue | Sequencing | Accession number | Source |
|---|---|---|---|---|
| IMR90 | H3K27ac | ChIP-seq | ENCSR002YRE | ENCODE |
| IMR90 | H3K4me3 | ChIP-seq | ENCSR087PFU | ENCODE |
| IMR90 | H3K4me1 | ChIP-seq | ENCSR831JSP | ENCODE |
| IMR90 | H3K36me3 | ChIP-seq | ENCSR437ORF | ENCODE |
| IMR90 | H3K27me3 | ChIP-seq | ENCSR431UUY | ENCODE |
| IMR90 | H3K9me3 | ChIP-seq | ENCSR055ZZY | ENCODE |
| IMR90 | input | ChIP-seq | ENCSR001BSB, ENCSR704GTT | ENCODE |
| IMR90 | RNA Polymerase II-Input | ChIP-seq | ENCSR000EFL | ENCODE |
| IMR90 | RNA Polymerase II | ChIP-seq | ENCSR000EFK | ENCODE |
| IMR90 | mRNA | RNA-Seq | ENCSR00CTQ | ENCODE |
| HepG2 | RNA Polymerase II-Input | ChIP-seq | ENCSR000EEM | ENCODE |
| HepG2 | RNA Polymerase II | ChIP-seq | ENCSR000EEN | ENCODE |
| HepG2 | H3K27ac, H3K4me3, H3K4me1, H3K36me3, H3K27me3, H3K9me3, Histone mark input, RNA-Seq | ChIP-seq | EGAD00001002527 | DEEP |
| K562 | mRNA | RNA-Seq | SRR315336, SRR315337 | European nucleotide archive |

**Table 1:** Experimental data used in this study. ChIP-seq data from 6 core histone modifications was used in this analysis.

### 2. Summary statistics

The summary statistics of transcription units predicted by EPIGENE, STRINGTIE and CUFFLINKS can be seen in Table 2,3 and 4.

| | genes | + strand | - strand | median length |
|---|---|---|---|---|
| **all** | 24,571 | 13,410 | 11,161 | 7,800 |
| **gencode V19 + chess 2.1 same strand overlap** | 18,184 | 9,774 | 8, 410 | 9,800 |
| **gencode V19 + chess 2.1 any overlap** | 23,542 | 12,921 | 10,621 | 8,400 |
| **no match** | 1,029 | 489 | 540 | 2,000 |

**Table 2:** Summary statistics of transcription units predicted by EPIGENE

| | genes | + strand | - strand | median length |
|---|---|---|---|---|
| **all** | 101,656 | 50,636 | 51,020 | 5,481 |
| **gencode V19 + chess 2.1 same strand overlap** | 93,006 | 46,448 | 46,558 | 6,719 |
| **gencode V19 + chess 2.1 any overlap** | 97,300 | 48,531 | 48,769 | 6,110 |
| **no match** | 4,356 | 2,105 | 2,251 | 613 |

**Table 3:** Summary statistics of transcription units predicted by STRINGTIE

| | genes | + strand | - strand | median length |
|---|---|---|---|---|
| **all** | 32,079 | 15,262 | 15,095 | 8,851 |
| **gencode V19 + chess 2.1 same strand overlap** | 26,452 | 12,986 | 12,671 | 16,486 |
| **gencode V19 + chess 2.1 any overlap** | 27,157 | 13,320 | 13,042 | 15,392 |
| **no match** | 4,992 | 1,942 | 2,053 | 962 |

**Table 4:** Summary statistics of transcription units predicted by CUFFLINKS

### 3. Robustness of the EPIGENE model

The robustness of EPIGENE was examined by testingK562-trained models on two other cell lines: IMR90 and HepG2. The similar performance score of independent EPIGENE models reflected its robustness.

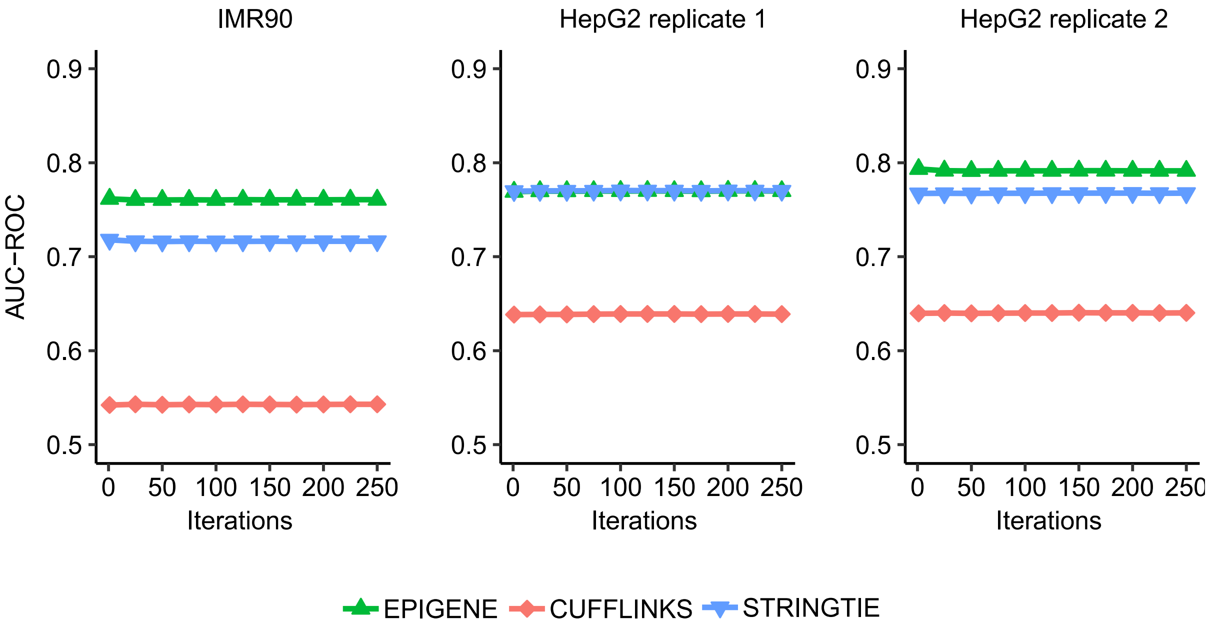

**Figure S1:** Comparison of K562 trained EPIGENE models with STRINGTIE and CUFFLINKS across 2 cell lines. ROC curves show that EPIGENE achieves a superior performance for 2 data sets compared to other approaches.

### 4. False positives due RNA-Seq mapping artefacts

We investigated the cause of higher AUC for EPIGENE compared to RNA-Seq based approaches and found that this due to slightly higher number of false positive resulting due to RNA-Seq mapping artefacts.

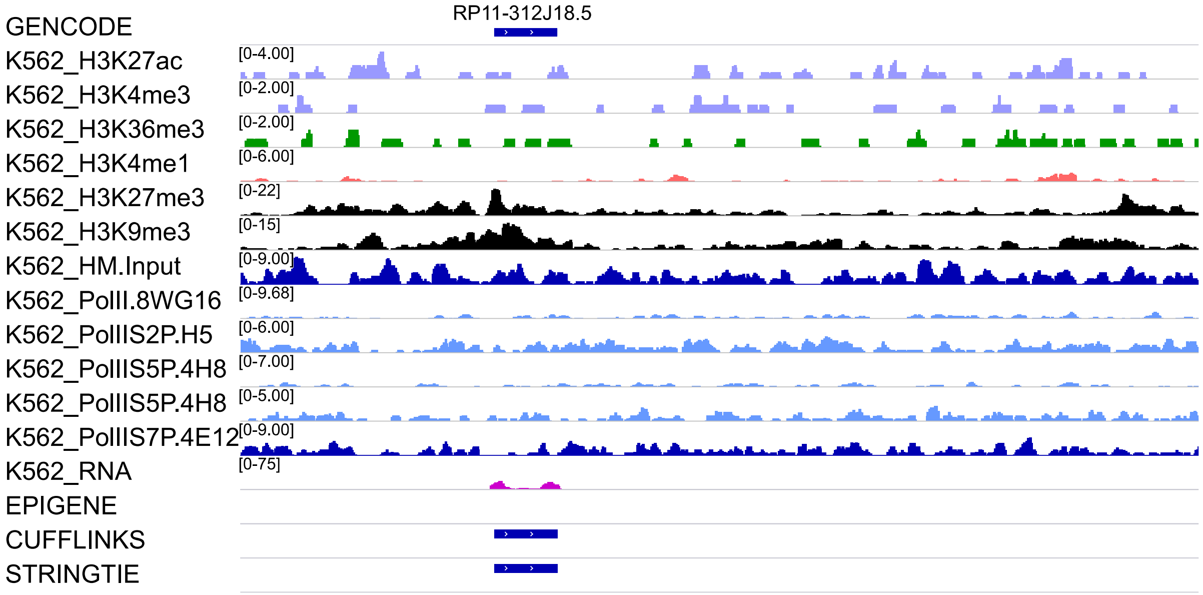


**Figure S2:** An example of CUFFLINKS and STRINGTIE due to spurious read mapping. This is a repetitive sequence occurring in chromosome 1,5,6,X, We observe an enrichment of repressive histone modifications like H3K27me3 and H3K9me3 (tracks shown in black) indicating that this is a repressive region.

### 5. Distribution of EPIGENE predictions across cell lines

We create a consensus TU set using the approach presented in section 4.7 of manuscript to identify cell specific TUs.

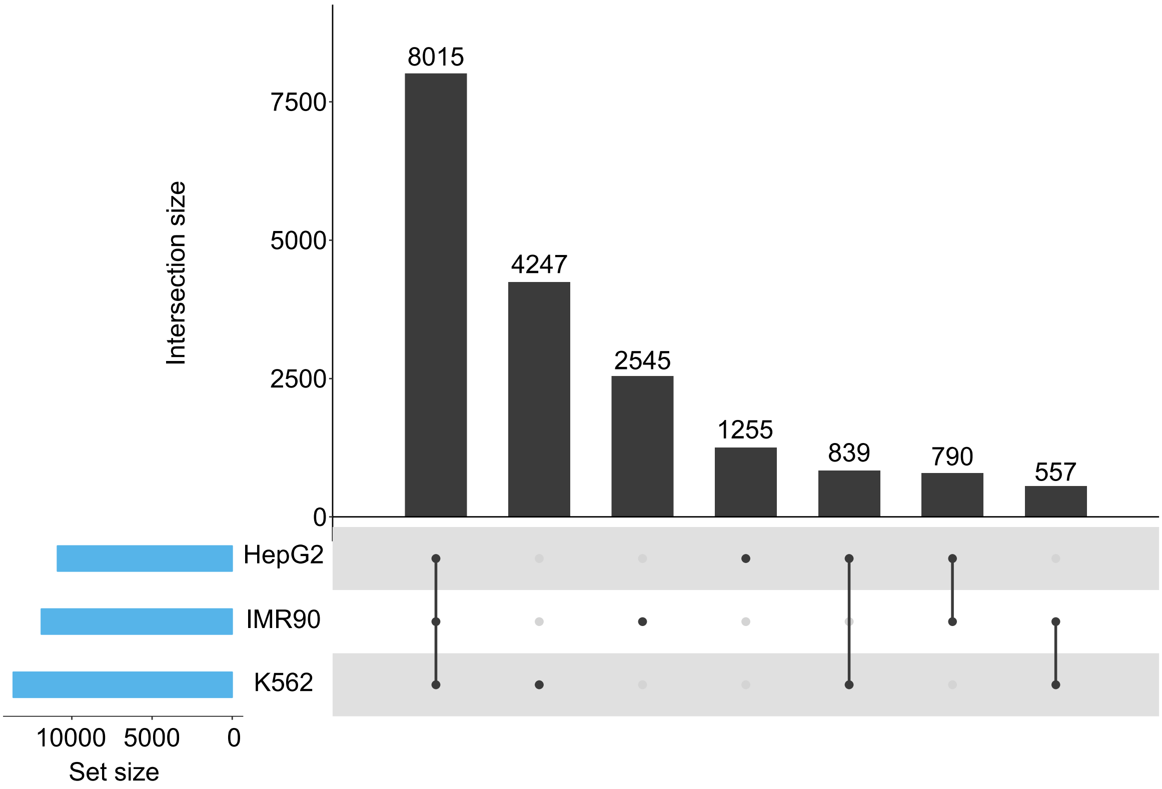


**Figure S3:** Distribution of EPIGENE TUs across cell lines

#### 6. Summary statistics of EPIGENE TUs overlapping miRNAs

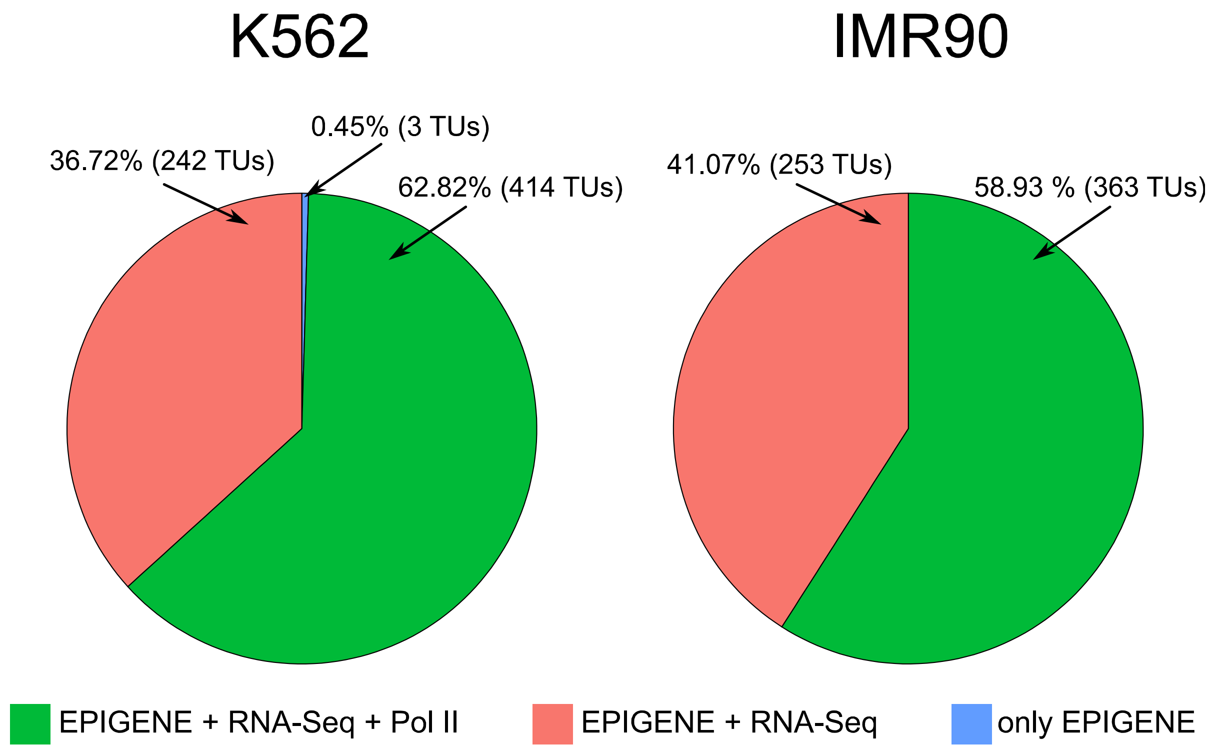


**Figure S3:** Majority of EPIGENE TUs overlapping miRNAs can be explained by RNA-Seq and Polymerase II evidence
